# Overcoming the skull barrier for noninvasive transcranial functional ultrasound imaging in marmosets

**DOI:** 10.64898/2026.03.03.708930

**Authors:** Hamidreza Ramezanpour, Amirhossein Asadian, Jeffrey Schall, Liya Ma

**Affiliations:** Department of Biology, York University, Toronto; Center for Vision Research, York University, Toronto; Department of Psychology, York University, Toronto; Centre for Integrative and Applied Neuroscience, Toronto

**Author notes:** **Correspondence to:** Hamidreza Ramezanpour, Department of Biology & Centre for Vision Research, York University, Toronto, Ontario, Canada, or Liya Ma, Department of Psychology & Centre for Vision Research, York University, Toronto, Ontario, Canada. These authors contributed equally to this work (Hamidreza Ramezanpour, Amirhossein Asadian).

**Keywords:** Functional Ultrasound Imaging, EDTA, Marmoset, Somatosensory Cortex, Isoflurane

## Abstract

Functional ultrasound (fUS) imaging enables high-resolution mapping of brain hemodynamics but is limited in primates by acoustic attenuation of the cranium. With topical application of ethylenediaminetetraacetic acid (EDTA), we were able to visualized cortical and deep vasculature in marmoset brains. We localized robust, stimulus-locked blood flow changes in cortical and subcortical regions contralateral to peripheral tactile stimulation. Beyond stimulus-evoked responses, resting-state cerebral blood flow also varied with the depth of isoflurane anesthesia. These findings advance translational ultrasound neuroimaging by offering a non-destructive alternative to a craniotomy.

## Introduction

Functional ultrasound (fUS) imaging offers an unparalleled combination of spatial and temporal resolution for mapping brain-wide hemodynamics (Macé et al., 2011), yet its application in large brains such as in nonhuman primates (NHPs) and humans is fundamentally constrained by skull-induced acoustic attenuation. In these species, fUS procedures require craniotomy (Dizeux et al., 2019; Blaize et al., 2020; Norman et al., 2021; Claron et al., 2023; Griggs et al., 2025; Liu et al., 2025) or skull thinning, which limit their longitudinal, translational and clinical applications. The only exception is in newborn babies, whose anterior fontanelle provides a natural acoustic window over part of the brain (Demene et al., 2017) for a limited time in early development.

Prior efforts to overcome the skull barrier have focused on hardware and computational strategies. These include adaptive focusing and beamforming with array transducers, CT-based correction (Aubry et al., 2003; Vignon et al., 2006), numerical and ray-based transcranial phase aberration correction (Wang et al., 2025a), and the use of lower ultrasound frequencies to reduce attenuation and scattering (Deffieux et al., 2018; Rabut et al., 2020). However, these methods typically trade spatial resolution for penetration depth or require complex calibration and modelling procedures.

By contrast, chemical modulation offers an orthogonal, potentially simpler route toward practical transcranial imaging. A recent study by Wang et al. (2025b) introduced a fundamentally different strategy by chemically modulating skull acoustic properties in mice. They demonstrated that topical application of ethylenediaminetetraacetic acid (EDTA), an FDA-approved calcium-chelating agent, transiently reduces skull mineral content, reducing acoustic impedance and sound speed toward values closer to soft tissue. In mice, this approach enabled near-lossless ultrasound transmission through the intact skull, restoring vascular signal intensity and spatial resolution to levels comparable to skull-removed preparations, and supporting reliable detection of stimulus-evoked fUS signals (Wang et al., 2025b).

It remains unclear whether this acoustic transparency strategy can be extended to non-human primates, whose skull curvature, laminar organization, and mineral composition more closely resemble those of humans. Here we test whether localized, well-controlled EDTA-mediated skull conditioning enables transcranial fUS imaging in the common marmoset, an important non-human primate model for systems and translational neuroscience (Mitchell and Leopold, 2015).

## Results

Local application of EDTA chelates divalent calcium ions within the skull, transiently reducing mineral content and reducing the acoustic impedance of the treated bone (Fig. 1A). Consequently, ultrasound wave propagation occurs with reduced reflection, scattering, and refraction at the bone interface (Fig. 1B). Hence, we applied EDTA treatment to the intact skull within a 3d-printed enclosure secured to the cranium of two marmosets (Fig. 1C). The treatment enabled stable fUS imaging of structural vascular contrast and stimulus-evoked hemodynamic responses.

**Figure 1.**
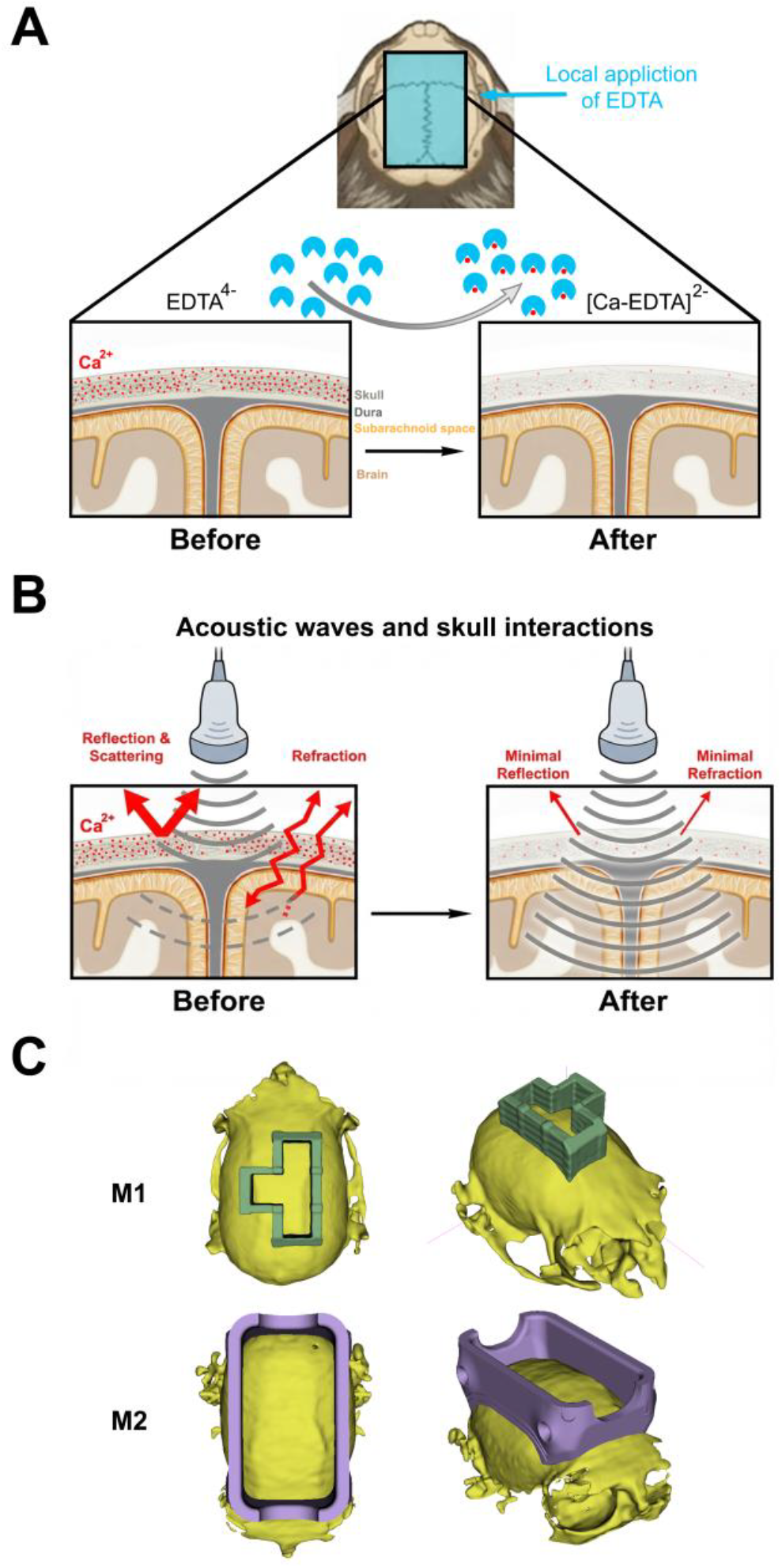
Experimental setup and EDTA-based skull treatment in marmosets. (A) EDTA chelates divalent calcium ions (Ca^2+^), forming [Ca–EDTA]^2-^ complexes and transiently reducing skull mineral content and acoustic impedance. (B) Prior to treatment, acoustic waves undergo reflection, scattering, and refraction at the mineralized bone interface. Following calcium chelation, impedance mismatch is reduced, resulting in more efficient transmission with minimal reflection and refraction. (C) Chambers secured to skull of marmoset 1 and 2 enabled topical application of EDTA. In Monkey 1, a T-shaped well was used to enable acquisition of both coronal and sagittal fUS imaging planes centered on the somatosensory cortex. In Monkey 2, a symmetric rectangular well was used, allowing uniform bilateral coverage of the targeted skull region.

### EDTA treatment enhances skull acoustic transparency and vascular visualization

Topical EDTA application produced a marked enhancement of skull acoustic transparency. After EDTA, two-dimensional fUS images showed increased signal clarity and finer vascular definition (Fig. 2A–C). Sagittal and coronal sections revealed robust visualization of the cortical vascular network, with the treated hemisphere consistently exhibiting higher image quality than the untreated side within the same animal (Fig. 2B). Although cortical vessels were clearly resolved following treatment, signal penetration into deeper subcortical structures remained comparatively limited (Fig. 2C).

**Figure 2.**
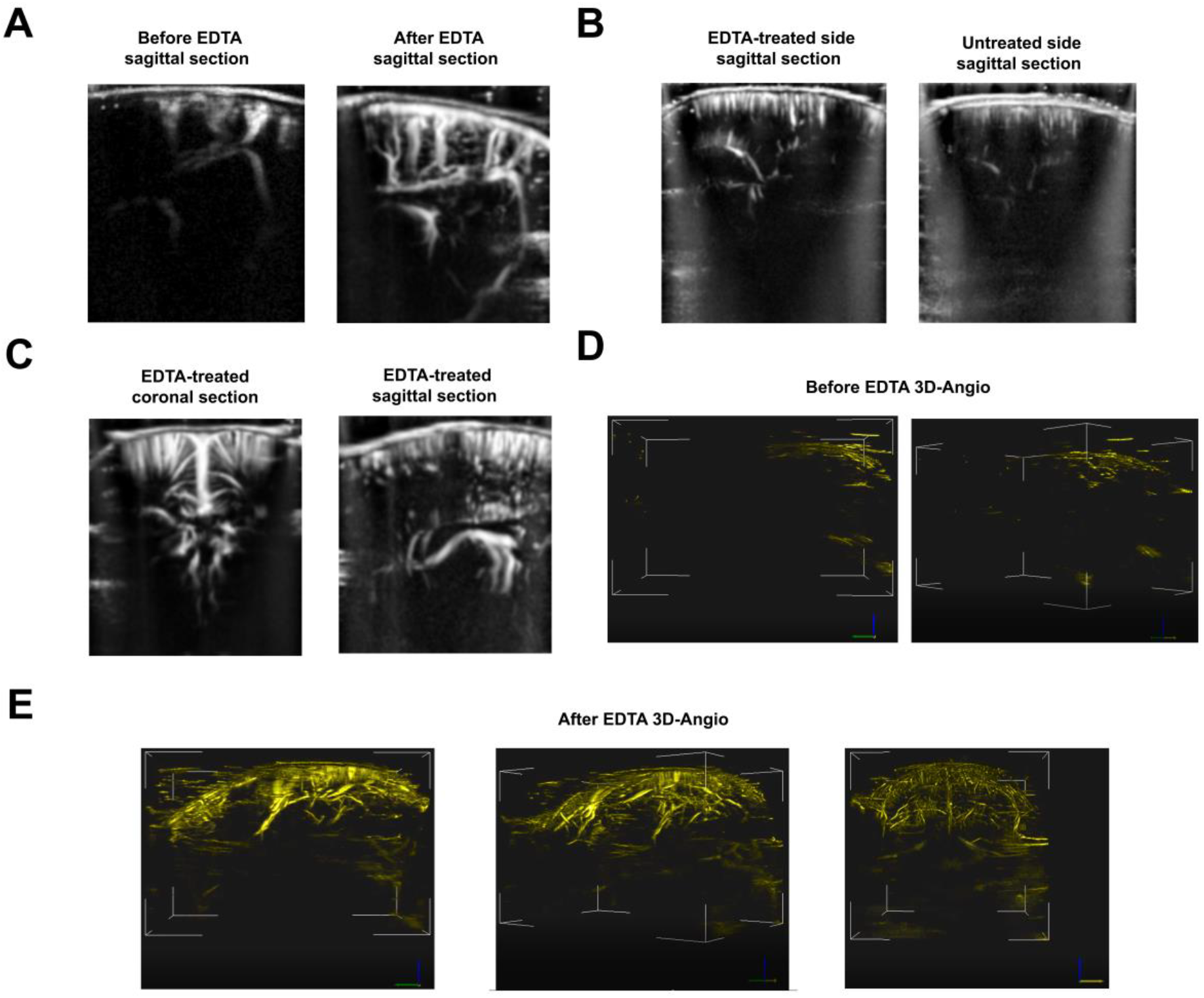
Effects of EDTA treatment on skull acoustic transparency and vascular visualization. (A) Representative sagittal scan before and after EDTA treatment in Monkey 1. (B) Example scan demonstrating improved image quality on the EDTA-treated side compared to the untreated hemisphere in Monkey 2. (C) Additional representative coronal and sagittal sections acquired after EDTA treatment in Monkey 1. (D) Three-dimensional whole-brain angiographic reconstruction acquired before EDTA treatment. (E) Three-dimensional angiographic reconstructions acquired after EDTA treatment.

The improved imaging quality was evident in three-dimensional whole-brain angiographic reconstructions. Before EDTA treatment, angiographic imaging was obscured by pronounced acoustic shadowing (Fig. 2D). After EDTA treatment, angiographic imaging revealed a dramatic enhancement in cortical vessel visualization across multiple viewing angles, along with partial recovery of signal from deeper vascular structures (Fig. 2E). These results demonstrate that localized EDTA application induces measurable changes in skull acoustic properties that translate into improved two- and three-dimensional ultrasound-based vascular imaging.

### Somatosensory cortical activation evoked by foot stimulation

We next mapped stimulus-evoked somatosensory responses in anesthetized marmosets. Peripheral tactile stimulation of the foot elicited robust, spatially localized increases in cerebral blood volume that were time-locked to stimulation blocks and lateralized to the hemisphere contralateral to stimulation (Fig. 3). These responses were consistently observed in both marmosets.

**Figure 3.**
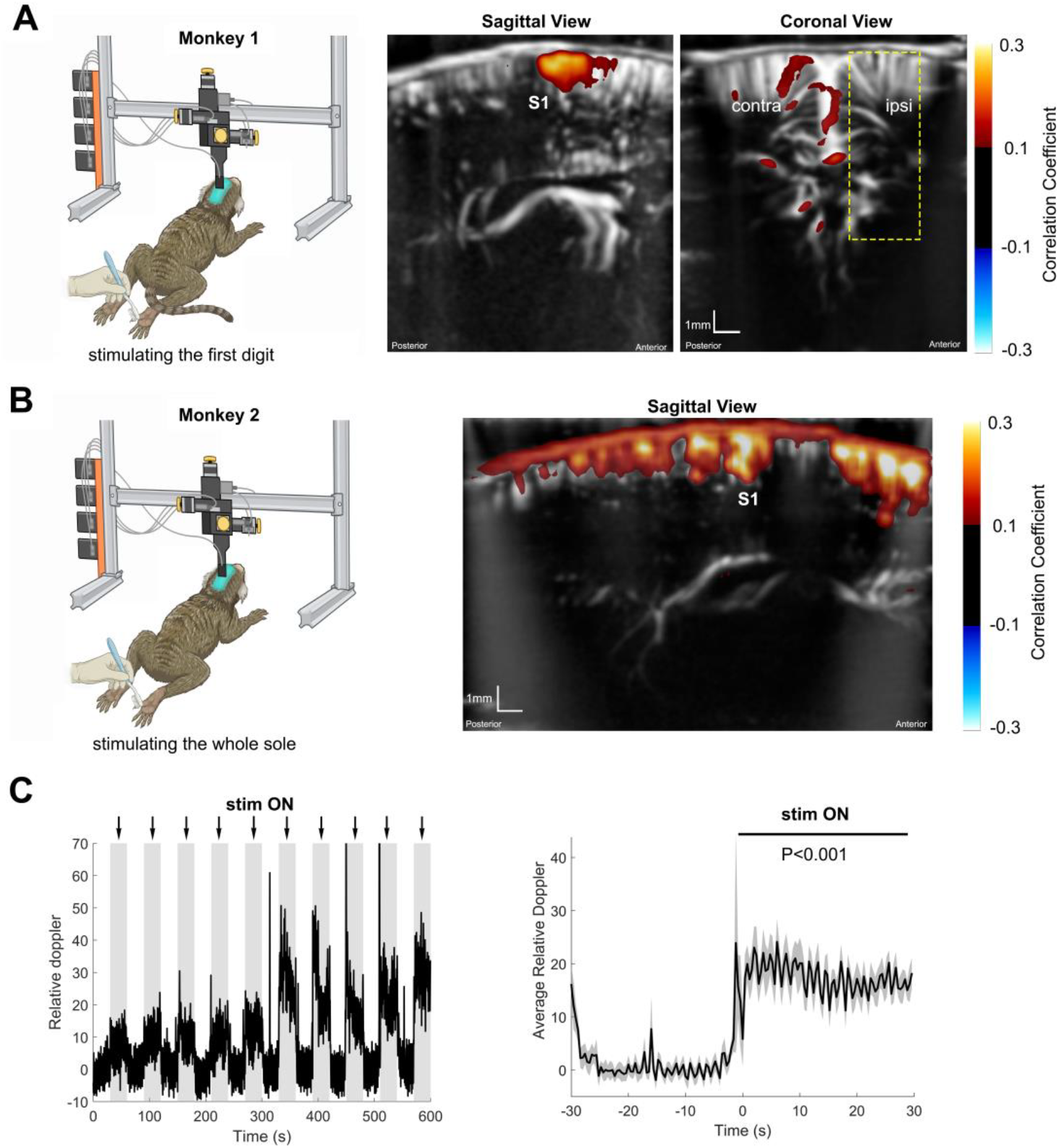
Somatosensory-evoked fUS responses to foot stimulation in anesthetized marmosets. (A) Monkey 1. The first digit was stimulated with a brush in 30 s ON / 30 s OFF blocks while fUS imaging was performed in sagittal and coronal planes targeting primary somatosensory cortex (S1). Robust stimulus-locked blood flow increases were observed in cortical regions consistent with S1, with responses confined to the contralateral hemisphere. Subcortical foci were also observed. (B) Monkey 2. Stimulation of the entire plantar surface of the foot elicited strong blood flow responses across multiple cortical regions following EDTA-mediated skull treatment, with maximal activation in S1, and additional responses in parietal and premotor cortices. (C) Stimulus-locked power Doppler signals extracted from an S1 region of interest, and the average response across 10 stimulation blocks, demonstrating a significant increase in blood flow following stimulation (p < 0.001, Wilcoxon signed-rank test).

In Monkey 1 (Fig. 3A), tactile stimulation of the first digit of the right hind foot was delivered manually using a brush in a block design consisting of 30 s stimulation ON and 30 s OFF, while fUS imaging was performed in both coronal and sagittal planes. Imaging planes were selected to target the primary somatosensory cortex (S1) based on marmoset MRI coordinates (Cléry et al., 2020). The coronal plane was positioned ∼6 mm anterior to the interaural plane, while the sagittal plane was placed ∼2 mm lateral to the midline, corresponding to the expected foot representation within S1 (Cléry et al., 2020). Robust stimulus-locked increases in cerebral blood flow were observed in cortical regions consistent with the expected S1 foot area, with strong activation evident in both sagittal and coronal views. Notably, in the coronal plane, activation was confined to the contralateral hemisphere, consistent with the known lateralization of somatosensory processing.

In addition to the prominent cortical responses, several subcortical activation foci were also observed. Based on their anatomical location, these signals may correspond to structures within the basal ganglia (e.g., caudate nucleus) and thalamic nuclei, but precise anatomical identification is not yet possible.

Results from Monkey 2 (Fig. 3B) further demonstrate the robustness of somatosensory responses following EDTA application. Tactile stimulation was applied to the entire plantar surface of the foot using a brush in the 30 s ON / 30 s OFF block design. This manipulation elicited strong and spatially extended CBV responses across multiple cortical regions, with the most pronounced activation observed in S1. Additional activation was detected in adjacent parietal cortex, likely corresponding to area 7, as well as in premotor/prefrontal regions, suggesting engagement of higher-order sensorimotor networks.

Quantitative analysis of the fUS signal is shown in Fig. 3C. Doppler power signals extracted from a region of interest encompassing S1 exhibited clear stimulus-locked responses, tightly time-locked to the stimulation epochs. Averaging the Doppler signal across 10 stimulation blocks revealed a robust increase in blood flow following stimulus onset, which was statistically significant (p < 0.001, Wilcoxon signed-rank test). These findings demonstrate reliable and reproducible somatosensory-evoked hemodynamic responses in marmoset cortex after EDTA treatment.

### Modulation of CBV by isoflurane anesthesia

All experiments in the present study were conducted in anesthetized marmosets. Because anesthesia is known to exert effects on cerebral hemodynamics, we examined how isoflurane depth modulated CBV signals. This was performed in one animal (M2) in a terminal experiment in which CBV responses were quantified across multiple cortical and subcortical regions during a stepwise increase in isoflurane concentration (Fig. 4). Systematic sampling of regional CBV responses was accomplished by defining cortical ROIs from anterior to posterior cortex at 1 mm intervals plus a subcortical ROI (Fig. 4A). Across ROIs, CBV exhibited a non-linear dependence on isoflurane concentration. Mean CBV increased from 2.0–2.5% to 3.5% isoflurane and then decreased at 4.5–5.0% (Fig. 4B). This inverted U-shape pattern was consistent across cortical regions, with individual ROIs showing similar dose– response trajectories. The time course of this modulation is shown in Fig. 4C. As anesthesia depth increased, CBV rose gradually, reached a maximum during intermediate isoflurane levels, and subsequently declined as isoflurane concentration increased further. Individual ROI traces closely followed the global mean trajectory, indicating a widespread effect of anesthesia depth on vascular dynamics.

**Figure 4.**
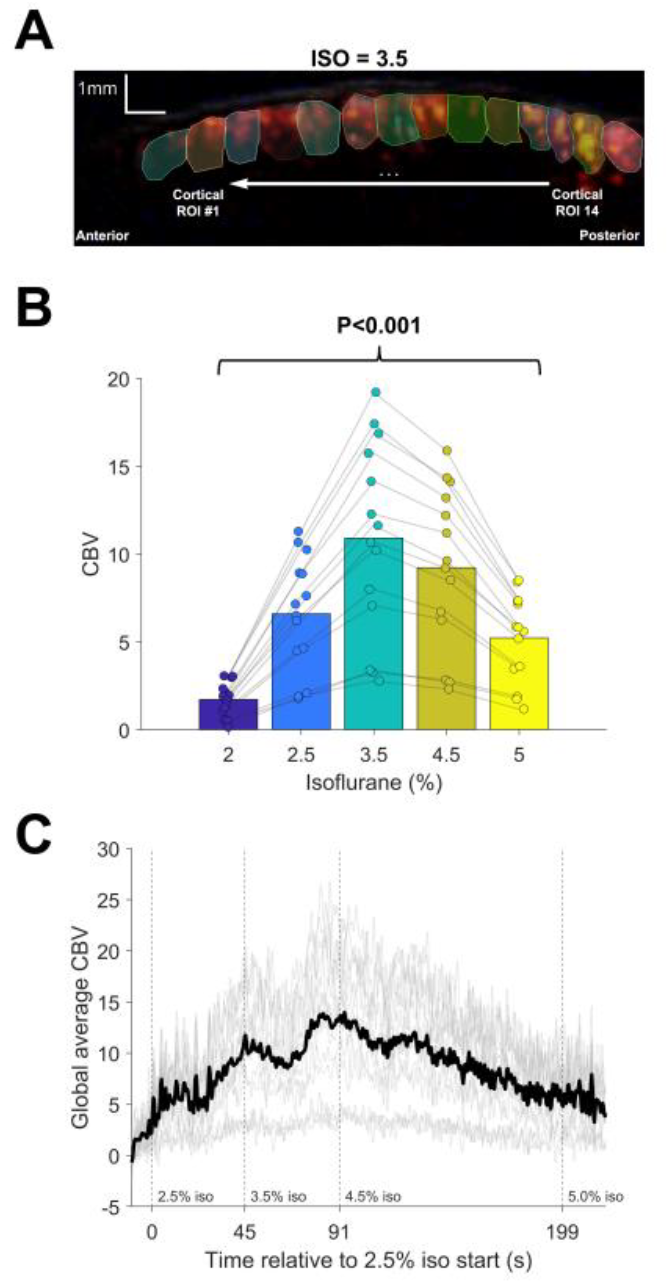
Isoflurane anesthesia induces a non-linear modulation of cerebral blood volume (CBV) across cortical regions. (A) Definition of regions of interest (ROIs) used for CBV analysis. Cortical ROIs were defined from anterior to posterior cortex at ∼1 mm spacing, together with a representative subcortical ROI, shown here overlaid on a sagittal fUS image acquired at 3.5% isoflurane. (B) Mean CBV values across ROIs plotted as a function of isoflurane concentration. Individual ROI values are plotted across doses, revealing a non-monotonic, inverted U-shaped relationship with a peak at intermediate anesthesia levels. (C) Time course of CBV during the stepwise isoflurane ramp. CBV time courses from individual ROIs (gray) and the global mean across ROIs (black) are aligned to the onset of 2.5% isoflurane. Vertical dashed lines indicate transitions between anesthesia levels, illustrating a rise in CBV at intermediate doses followed by a decline at higher concentrations.

## Discussion

### Functional specificity enabled by EDTA-mediated transcranial fUS

The somatosensory responses observed here validate the functional specificity enabled by EDTA-mediated skull conditioning. Tactile stimulation of the foot elicited robust, stimulus-locked increases in cerebral blood volume that were lateralized to the hemisphere contralateral to stimulation and consistent across coronal and sagittal imaging planes. These features closely mirror findings from fMRI studies in primates, in which contralateral S1 and associated subcortical nuclei exhibit strong, stimulus-locked hemodynamic responses to limb stimulation (Cléry et al., 2020). In addition to cortical activation, we observed several subcortical foci that, based on their anatomical location, may correspond to thalamic and basal ganglia structures within sensorimotor circuits. At this stage, however, fUS images have not been registered to a CT–MRI marmoset brain atlas, and therefore regional assignments remain provisional. Ongoing work aims to integrate atlas-based registration with transcranial fUS imaging to enable more definitive localization in future studies. Together, these findings indicate that EDTA-enabled transcranial fUS preserves functional fidelity while substantially reducing invasiveness.

Prior work has shown that topical EDTA treatment can reversibly increase skull acoustic transparency in rodents and enhance ultrasound transmission through ex vivo human skull samples (Wang et al., 2025b). Extending this approach to marmosets, a non-human primate model, bridges a critical anatomical and biophysical gap between rodent preparations and human applications. The recovery of cortical vascular signals and robust stimulus-evoked functional responses through the intact primate skull therefore represents an important translational step toward minimally invasive, ultrasound-based functional imaging (Martinez de Paz and Macé, 2021) and, potentially, neuromodulation in humans (Darmani et al., 2022).

### Non-linear modulation of CBV by anesthesia depth and relation to prior fMRI work

The observed inverted U-shaped relationship between cerebral blood volume (CBV) and isoflurane concentration, consistent across multiple brain regions, arises from the competing influences of isoflurane’s direct vasodilatory effects on cerebral vasculature and its dose-dependent inhibition of neuronal excitability and neurovascular coupling. At low to intermediate isoflurane concentrations, the agent functions as a potent cerebral vasodilator, thereby increasing baseline perfusion and elevating CBV. Concurrently, isoflurane augments inhibitory neurotransmission through potentiation of γ-aminobutyric acid type A (GABAA) receptors (Masamoto et al., 2009; Tsurugizawa et al., 2016). As anesthetic depth increases, these neuronal suppressive effects predominate, leading to substantial decreases in neuronal discharge rates, synaptic transmission, and cerebral metabolic demand (Tsurugizawa et al., 2016). Consequently, the resultant hemodynamic response exhibits a non-monotonic profile, peaking at intermediate anesthetic levels and attenuating at higher concentrations, in alignment with patterns observed across widespread cortical ROIs in Fig. 4.

Analogous dose-dependent, non-linear modulations by isoflurane have been documented in rodent functional magnetic resonance imaging (fMRI) investigations. In rats and mice, baseline blood-oxygen-level-dependent (BOLD) signals and resting-state functional connectivity demonstrate inverted U-shaped or biphasic dependencies on isoflurane dosage, characterized by peak signal amplitude and variance at intermediate concentrations and attenuation under deeper anesthesia (Tsurugizawa et al., 2016). More broadly, prior work has emphasized that anesthetic depth profoundly alters both baseline and evoked hemodynamic responses, rather than merely scaling neuroimaging metrics as a uniform gain modulator (Aksenov et al., 2015). Our results extend these observations to a CBV-based, high-spatiotemporal-resolution imaging modality in a non-human primate model and underscore that anesthesia level should be treated as a critical experimental factor, and explicitly reported, in fUS studies.

### Future directions

Several important directions follow naturally from this work. First, systematic exploration of longer EDTA application durations may further enhance acoustic transparency and improve access to deeper cortical and subcortical structures, but will require careful characterization of safety, reversibility, and tissue tolerance in primates. Second, integration of transcranial fUS data with CT–MRI–based marmoset brain atlases will be essential for precise anatomical localization and for resolving subcortical activation patterns. Finally, combining EDTA-enabled transcranial fUS with ultrasound localization microscopy (ULM) using microbubbles offers the opportunity to achieve microvascular-resolution imaging in primates. Extending this approach to awake, behaving animals performing cognitive, motor, or visual tasks would enable circuit-level investigations with spatiotemporal precision that is currently inaccessible using existing non-invasive methods.

### Limitations and translational considerations

Several limitations should be considered when interpreting these findings. Although marmosets provide an important intermediate model, their skull properties remain distinct from those of adult humans, and EDTA-induced acoustic enhancement may not scale directly. Moreover, the current approach relies on topical EDTA application to the exposed skull and is not intended as a clinical procedure; long-term effects on bone integrity and surrounding tissues would require extensive evaluation. Finally, the field of view of current fUS probes remains limited for large brains, suggesting that near-term translational relevance may be confined to cortical targets. Nonetheless, this work provides a critical proof of principle motivating the development of clinically compatible, non-destructive strategies for improving transcranial ultrasound access in individuals with suboptimal acoustic windows. Together, these results establish EDTA-based skull conditioning as a viable strategy for enabling high-resolution transcranial fUS imaging in primates, bridging a critical gap between rodent studies and future human applications.

## Acknowledgment

We are grateful to Yaoheng (Mack) Yang and Zikay Wang for helpful discussions regarding the EDTA protocol. This work was supported by the Canada Foundation for Innovation (J.D.S.); York University’s Catalyzing Interdisciplinary Research Clusters program (J.D.S.); CFREF VISTA initiative (J.D.S., L.M.); CFREF Connected Minds initiative (L.M., J.D.S.); NSERC Discovery Grants (J.D.S., L.M.); and the Canada Research Chairs Program (J.D.S., L.M.).

## Methods

Two male adult common marmosets (*Callithrix jacchus*), both 4 years old, were used in this study. Animals were pair-housed in a temperature- and humidity-controlled facility under a 12-h light/dark cycle with ad libitum access to food and water. Environmental enrichment was provided according to institutional guidelines. All procedures, including anesthesia, surgical preparation, and imaging, were performed under isoflurane anesthesia with continuous physiological monitoring (heart rate, respiratory rate, and peripheral oxygen saturation) and in accordance with the guidelines of the Canadian Council on Animal Care on the use of laboratory animals and were also approved by the York University Animal Care Committee.

### fUS imaging

Imaging was performed under sterile conditions with animals positioned in a stereotaxic frame. Body temperature was maintained using a feedback-controlled heating pad. For pre-EDTA imaging, hair was removed, and acoustic coupling gel was applied directly to the scalp. fUS acquisitions were obtained using an Iconeus One system (Iconeus, Paris, France) with linear probes. For M1 imaging, an IcoPrime probe was used (center frequency 15 MHz; 128 elements; probe size 25 × 17.5 × 6 mm; field of view 14.1 mm; in-plane resolution ≈ 100 × 100 µm^2^). For M2 imaging, an IcoPrime-XL probe was used (center frequency 15 MHz; 192 elements; probe size 32 × 25.6 × 6 mm; field of view 21.1 mm; in-plane resolution ≈ 100 × 100 µm^2^).

For three-dimensional angiographic imaging, the probe was mounted on a motorized multi-axis linear translation stage (SLC-1740, SmarAct GmbH) and stepped sequentially along the sagittal axis in increments of 0.1 mm. At each position, 2D power Doppler images were acquired and subsequently combined to reconstruct a 3D angiographic volume.

Raw fUS data were exported in NIfTI format using IcoStudio software (Iconeus, Paris, France) and processed in MATLAB (R2022b, The MathWorks, Natick, MA). Data were converted to single precision, log-compressed, and temporally denoised by computing a median across frames within each acquisition block. Each slice was normalized to unit dynamic range, percentile-clipped to suppress outliers, and contrast-enhanced using adaptive histogram equalization (CLAHE, adapthisteq in MATLAB), followed by light Gaussian smoothing. The resulting images were upsampled with bicubic interpolation and displayed in grayscale.

### EDTA preparation and skull application

To transiently increase skull acoustic transparency, a 20% (w/v) EDTA solution was prepared following the same formulation and handling procedures described by Wang et al., with no modifications to the underlying chemical protocol (Wang et al., 2025b). EDTA powder was gradually dissolved in double-distilled water. Because EDTA exhibits limited solubility at near-neutral pH, sodium hydroxide (NaOH) was added incrementally under continuous stirring to facilitate dissolution. In some preparations, achieving a fully clear solution required the addition of slightly higher amounts of NaOH, which transiently increased the solution pH to approximately 8.0. In these cases, the pH was subsequently adjusted downward to physiological pH (7.4) by adding a small volume of hydrochloric acid (HCl) at the final step. Maintaining physiological pH was critical to minimize potential tissue irritation and to ensure safe topical exposure of the skull. After complete dissolution and pH adjustment, the solution was brought to the final volume and sterilized.

For marmoset experiments, EDTA was applied topically to the intact skull using a custom 3D-printed well-based approach (Fig. 1C). Following scalp incision and exposure of the skull surface, custom 3D-printed wells were temporarily affixed to the skull using a low-toxicity silicone adhesive (Kwik-Sil, World Precision Instruments, Sarasota, FL, USA), forming a sealed reservoir over the region of interest and allowing controlled containment of the EDTA solution.

The well was filled with freshly prepared EDTA solution and maintained for a total duration of 30–45 minutes. To ensure consistent chelation efficacy, the EDTA solution was replenished every 5 minutes by gently suctioning the existing liquid from the well and immediately replacing it with fresh solution. Upon completion of the treatment period, the EDTA was removed, and the skull surface was thoroughly rinsed with sterile saline prior to fUS imaging. Structural effects of EDTA treatment on the skull, including pre- and post-treatment changes in thickness and vascular visibility, are shown in Fig. 2.

Well geometry was adapted based on experimental goals. In Monkey 1, a T-shaped well was used to enable acquisition of both coronal and sagittal imaging planes centered on the somatosensory cortex (Fig. 1, Monkey M1, top view). In this animal, EDTA treatment was applied unilaterally, with the contralateral skull left untreated for within-subject comparison. In Monkey 2, a symmetric rectangular well was used (Fig. 1C, lower), allowing bilateral and uniform EDTA exposure over the targeted skull region. In this animal, prior to EDTA application, the skull surface was gently prepared using a dental drill. This step served two purposes: (1) removal of any residual silicone adhesive from previous procedures, and (2) mild thinning of the outer skull layer to facilitate EDTA penetration and improve chelation efficiency. Care was taken to avoid excessive thinning or thermal damage, and the skull surface was irrigated as needed during preparation.

### Activation map generation

Activation maps were generated using the Activation Map module in IcoStudio (version 2.3.0; Iconeus, Paris, France) which computes voxel-wise Pearson correlation coefficients between the Power Doppler time series and a binary stimulation pattern. The stimulation paradigm (30 s ON / 30 s OFF blocks) was entered using the preset interface. No hemodynamic response function convolution was applied. A correlation threshold of r ≥ 0.1 was used for visualization. The same correlation threshold and display scaling parameters were used across animals. For the purpose of better visualization, the extracted activations maps were overlaid on the contrast-enhanced images.

### ROI definition and CBV analysis across isoflurane levels

To examine the effect of anesthesia depth on cerebral blood volume (CBV), regions of interest (ROIs) were defined along the cortical surface and within a representative subcortical structure using fUS images. Cortical ROIs were placed sequentially from anterior to posterior cortex at approximately 1 mm spacing, enabling sampling of CBV responses across a broad extent of cortex. This analysis was performed in one animal (Monkey 2; M2). For each ROI, CBV time courses were extracted from the power Doppler signal.

CBV signals were temporally trimmed to include data from 10 s prior to the onset of the 2.5% isoflurane step through the end of the anesthesia ramp. The resulting time series were low-pass filtered using a zero-phase filter, with the cutoff frequency constrained relative to the sampling rate to avoid aliasing. Mean CBV values were then computed within predefined time windows corresponding to steady-state periods at 2.0% (baseline), 2.5%, 3.5%, 4.5%, and 5.0% isoflurane. The baseline condition was defined as the final 10 s immediately preceding the onset of 2.5% isoflurane.

For visualization, mean CBV values were displayed as bar plots with individual ROI values overlaid and connected across anesthesia levels (Fig. 4B). Temporal dynamics were visualized by aligning CBV time courses to the onset of the 2.5% isoflurane step (Fig. 4C).

## Notes

### Competing Interest Statement

The authors have declared no competing interest.

### Summary of Updates

This version of the manuscript has been revised to update the references.

